# Molecular stripping in the *NFκB/IκB/DNA* genetic regulatory network

**DOI:** 10.1101/027912

**Authors:** Davit A Potoyan, Weihua Zheng, Elizabeth Komives, Peter G Wolynes

**Affiliations:** Center for Theoretical Biological Physics and Department of Chemistry, Rice University, Houston TX 77005; Department of Chemistry and Biochemistry, University of California, San Diego CA

## Abstract

Genetic switches based on the *NFκB/IκB/DNA* system are master regulators of an array of cellular responses. Recent kinetic experiments have shown that *IκB* can actively remove NF*κ*B bound to its genetic sites via a process called ”molecular stripping”. This allows the *NFκB/IκB/DNA* switch to function under kinetic control rather than the thermodynamic control contemplated in the traditional models of gene switches. Using molecular dynamics simulations of coarse grained predictive energy landscape models for the constituent proteins by themselves and interacting with the DNA we explore the functional motions of the transcription factor *NFκB* and its various binary and ternary complexes with DNA and the inhibitor I*κ*B. These studies show that the function of the *NFκB/IκB/DNA* genetic switch is realized via an allosteric mechanism. Molecular stripping occurs through the activation of a domain twist mode by the binding of *IκB* which occurs through conformational selection. Free energy calculations for DNA binding show that the binding of *IκB* not only results in a significant decrease of the affinity of the transcription factor for the DNA but also kinetically speeds DNA release. Projections of the free energy onto various reaction coordinates reveal the structural details of the stripping pathways.

---

The binding and release of protein transcription factors from *DNA* are fundamental molecular processes by which genes are regulated in the cell. The pioneering studies of Jacob, Monod, Ptashne and Gilbert explained how these two processes, seeming inverses of each other, while being maintained in local chemical equilibrium, could still lead to robust genetic switches by coupling to protein synthesis and degradation which are kinetically controlled far from equilibrium processes (*1–4*). This classic picture, with the law of mass action at its core (*5,6*), suggests that understanding the molecular mechanism of the binding and release of transcription factors is of secondary interest compared with understanding the thermodynamics of protein-DNA recognition. The recent discovery of protein induced release of a *DNA*-bound transcription factor in the *NFκB/IκB/DNA* genetic switch changes this picture (*7*). The induced process, called ”molecular stripping”, opens up the possibility of molecular kinetic control of binding and release thus overturning the classical paradigm based only on thermodynamic control. In this paper, we use molecular dynamics simulations of coarse-grained but predictive energy landscape models of the proteins along with their interacting *DNA* to explore first how the *NFκB* transcription factor binds individually both to *DNA* and to its inhibitor *IκB* and then to study how an approaching *IκB* can strip the *NFκB* from a *DNA* molecule to which it has already been bound, by forming an intermediate ternary complex. These simulations show that each of the binary binding events involves conformational selection of different *NFκB* global conformations. Molecular stripping then occurs through an allosteric transition between these conformations that is induced by forming the ternary complex. Stripping involves changes in the patterns of local frustration in the proteins and an interchange of electrostatic interactions between the protein-protein and the protein-*DNA* partners. The simulations highlight the breaking of symmetry in the *NFκB* hetero-dimer complex with *IκB* as well as the important role of the PEST sequence of *IκB in the stripping.*

## Systems biology of *NFκB/IκB/DNA*

The transcription factor *NFκB* is central to the regulation of inflammatory responses, immune responses to infections and programmed cell death. *NFκB* is a collective name for a family of homo- and hetero-dimeric proteins which serve as a bridge from the extracellular signals to gene expression (*8*). The *NFκB* hetero-dimer p50p65 is the most abundant form, although other forms are also present in the cell. *NFκB* is activated by a diverse set of extracellular stimuli that through an enzyme cascade mark the *IκB* moiety for degradation subsequently freeing *NFκB* from its inhibited complex. In its active unbound form *NFκB* translocates to the nucleus thereby initiating the transcription of hundreds of genes. Due to its wide range of influence *NFκB* is considered a ”Master regulator switch” (*9*). Among the target genes of *NFκB* is its own inhibitor *IκBα* which we will refer to simply as *IκB*. Targeting *IκB* synthesis results in a negative feedback loop that governs the nuclear concentration of active *NFκB* leading to pulses of activation and inactivation (*10,11*). *IκB* strips the *NFκB* off from its *DNA* targets and masks the nuclear localization signal of the *NFκB* (*7*). The resulting tightly bound *NFκB*−*IκB* complex (*12*) is thus prevented from reentering the nucleus (*13,14*) thereby terminating all the associated genetic activities of *NFκB* to restore the cellular resting state (Fig. 1A)

**Figure 1:**
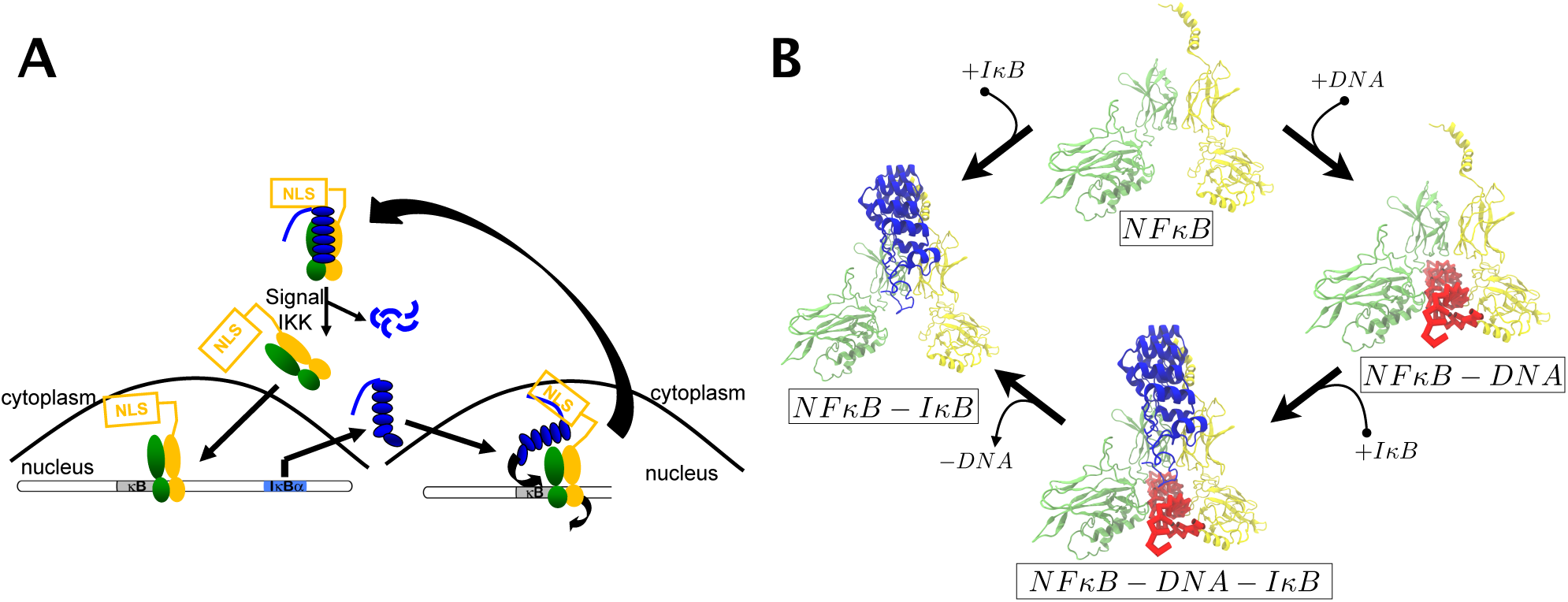
(A) Schematic systems biology view of *NFκB* regulation by its inhibitor *IκB*. Shown are the key steps of a negative feedback cycle which includes the *NFκB* activation by external stimuli, followed by DNA binding, *IκB* synthesis and molecular stripping of *NFκB* by *IκB* in conjunction with direct binding to free *NFκB* (B) Diagram showing the predicted structures used in the simulations. The names of complexes are shown at the bottom of the structures. Arrows indicate the thermodynamically favored directions for interconversion between the complexes given sufficient concentrations of the *IκB* and the DNA binding sites.

The most complete mathematical models of *NFκB* signalling have assumed fast local equilibria for *NFκ*B binding to the DNA (*15, 16*). Recent *in vitro* kinetic measurements of *IκB* binding to the complex of *NFκB* with DNA challenge this equilibrium view. *NFκB* dissociates from DNA in an *IκB* concentration dependent manner (*7,17*). Including this enhancement in models requires going beyond the purely equilibrium picture that envisions fast dissociation/association and that ignores competing binding at other gene sites. A revised systems biology scheme showing the central role of molecular stripping is described in Fig. 1A. *In vivo* there are several thousand *κB* binding sites (*18*), thus if dissociation from these target and decoy sites would not be fast enough there would be wasteful over-expression of genes. Rapid molecular stripping of *NFκB* from its DNA results in efficient down regulation of the *NFκB* targets ensuring the optimal expenditure of cellular resources. Our simulations demonstrate a feasible mechanism for this molecular stripping reaction.

## Structures, dynamics and thermodynamics

Structures of the p50p65 hetero-dimer of *NFκB* bound with the DNA and with its inhibitor *IκB* have already been resolved with X-ray crystallography (*14,19,20*). Unfortunately each of the available structures lacks at least one mechanistically crucial part of the *NFκB* molecule. The structure of *NFκB* − *DNA* (PDB ID: 1LE5) by Berkowitz et al (*19*) lacks the crucial nuclear localization signal (NLS) while the entire p50 N-terminal binding domain is missing from the two existing crystal structures (*14, 20*) for *NFκB* − *IκB* (PDB ID: 1IKN, 1NFI). To understand the binding and stripping mechanisms, more complete structures are needed. To construct these structures (Fig. 1B) we carried out simulations with a coarse-grained protein-DNA force field. We added the missing molecular fragments and re-generated the full structures of the various complexes by using the **A**ssociative memory, **W**ater mediated, **S**tructure and **E**nergy **M**odel (AWSEM) for the interactions within and among the proteins (*21*) and the 3 Sites Per Nucleotide (3SPN) model for the DNA (*22,23*). DNA and protein are coupled through steric exclusion and electrostatics modeled at the mean field level. AWSEM is a coarse grained predictive model based on the energy landscape formulation of protein folding theory (*24*). It accounts explicitly for physically motivated interactions such as hydrogen bonding and solvent interactions to model tertiary structure relationships, and incorporates ”evolutionary forces” via bioinformatic terms that act locally in sequence. Both kinds of interactions effectively sculpt the landscapes of naturally occurring proteins to be highly funneled towards the native state. AWSEM has proved successful in predicting structures of both protein monomers (*21*) and dimers (*25*). The 3SPN force field is a coarse grained model for the DNA that also includes physically motivated terms, such as base pairing, base stacking and electrostatic repulsion of phosphate groups which give rise to the double helical structure of the DNA with appropriate thermal and mechanical properties (*26, 27*). It gives a good account of the range of structural diversity of double helical DNA (*26*). Details of the models can be found in the Supplementary Materials section.

The three dimensional structures predicted by our code compare well with the available crystal structures of *NFκB* − *DNA*. The tertiary and secondary levels of structure are reproduced in the simulations with high fidelity. The binding interface between protein and DNA is also well predicted. We also can compare the predicted complete structure of the complex *NFκB* − *IκB* with the crystal structure of an *NFκB* − *IκB* complex (PDB ID: 1IKN) in which the p50 binding domain has been removed (Fig. S1). The crystal structure of the truncated complex aligns well with the simulated structure in the dimerization domain region. In the crystal structure of the truncated form, however, the p65 N-terminal binding domain has twisted further from its location in the *NFκB* − *DNA* complex than it has in the predicted complete structure. This larger domain twist in the artificially truncated crystal structure results from the higher flexibility of the p65 N-terminal domain allowed by the absence of steric conflicts with the neighboring p50 N-terminal domain (Fig. S1). When both N-terminal domains are present, the orientation of the domains is sterically forced to be closer to the orientation seen in the crystal structure of *NFκB* − *DNA*. Simulations of a hetero-dimer in which the p50 N-terminal binding domain has been removed concur in predicting this additional twist. In the following sections we make use of our predicted full structures of the bound complexes to follow their formation dynamics and reaction pathways.

Naturally occurring proteins are minimally frustrated as a result of evolutionary development. The ability of proteins to fold into unique native configurations on reasonable time-scales is successfully explained by folding theories resting on the principle of minimal frustration (*28–30*). Frustration is not completely eliminated via evolutionary selection because functional constraints also sculpt the energy landscapes to allow functional motions of the folded proteins in the folded states and binding reactions (*30, 31*).The folded protein ensemble occupies the dominant basin of the free energy landscape where frustration properly localized in key regions of the protein chain opens up routes for specific functional dynamics.

Frustration in biomolecules can be readily quantified and visualized using a spatially local version of the principle of minimal frustration (*30*). An algorithm to do this is implemented in the Frustratometer (*32*) which estimates the energetic contribution of contacts to the stability of the native state and compares these energies in the native structure to the distribution of energies that would be obtained by performing residue substitutions in the local environment. Native contacts whose energies are sufficiently deep in the energetically stabilizing part of the distribution are called minimally frustrated and are drawn as green lines on the structure while the native contacts that are found sufficiently far in the energetically destabilizing region are called frustrated and are drawn as red lines. The quantitative details of this algorithm including electrostatic effects have been discussed in previous papers (*30, 32, 33*).

The frustration patterns of the *NFκB* were analyzed in the structure of free *NFκB*, *NFκB* bound with the *DNA* and *NFκB* bound with *IκB*. When electrostatic effects are included, localized high levels of frustration are found in the binding pocket of the *NFκB* dimer when considered by itself (Fig. 2A) or when the protruding PEST sequence of the *IκB* is truncated in the *NFκB* − *IκB* dimer (Fig. 2B). This frustration is largely caused by the positively charged residues which are located in close proximity to each other. Binding DNA or the *IκB* PEST sequence both of which are negatively charged mitigates this frustration so that the necklace of frustration in the binding pocket (Fig. 2A-B) becomes unfrustrated after DNA binds (Fig. 2C) or when the PEST sequence of the *IκB* moves into the pocket (Fig. 2D). Thus frustration analysis already hints that binding *IκB* with its PEST can facilitate the stripping of *NFκB* from DNA by relieving frustration in the free *NFκB* that forms if the DNA dissociates by itself.

**Figure 2:**
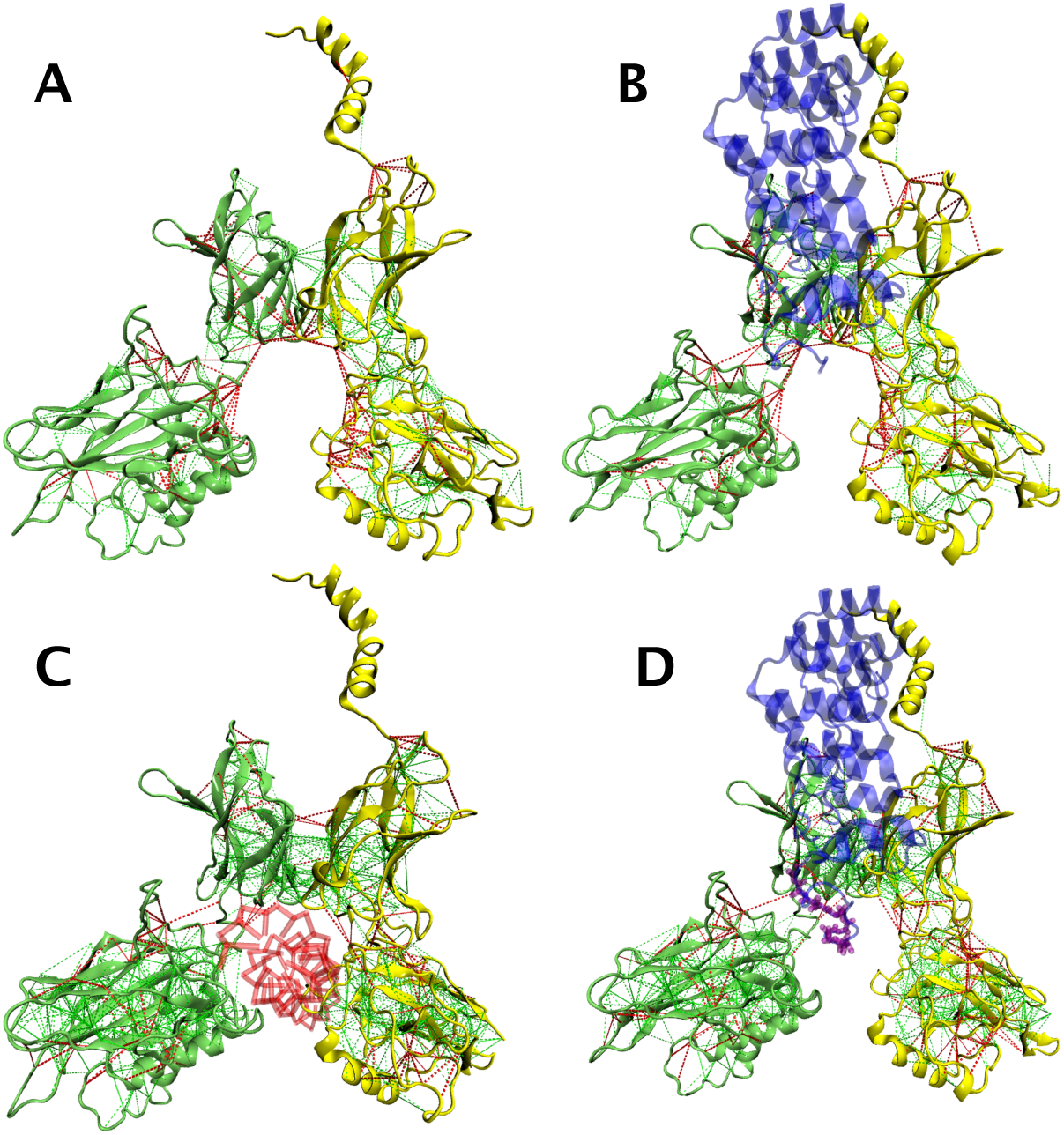
Frustration analysis with electrostatics for the bound and free forms of the *NFκB*. Shown are the frustration patters in (A) unbound NFkB dimer (B) *NFκB* − Iκ*B* complex with the PEST sequence removed (C) *NFκB* − DN*A* complex (D) *NFκB* − *IκB complex.*

This origin of stripping is underlined by free energy calculations using umbrella sampling simulations of *DNA* binding to *NFκB* by itself and of DNA binding to a pre-existing *IκB* : *NFκB* complex. The starting structures for these simulations are the predicted structures previously described as well as a model of the ternary complex of *IκB* − *NFκB* − *DNA* also created by AWSEM. From these starting points we used umbrella sampling to sample more broadly the complete ensemble of conformational states intermediate between those where the partners are fully bound and other conformations where they are completely released. These simulations capture the steps of molecular stripping.

The distances between either the center of mass of the dimerization domains with the center of mass of the *DNA* or with the center of mass of the *IκB* were used as harmonic biasing terms for the umbrella sampling. While the *in vitro* experiments were carried out on the short piece of *DNA* which we used in simulations (Fig. S2), we also carried out simulations that better mimic the way the transcription factor would approach a much longer full length genomic *DNA in vivo*, by constraining the orientation of the *DNA* to remain parallel to the orientation of the initial bound state. Without this constraint an intermediate forms in which the protein complex binds to the apex of the short *DNA* fragment (Fig. S3). Introducing the orientational constraint removes this artificial effect of the finite length of the *DNA* fragment studied *in vitro*.

The 1D Free energy profiles for the biologically more relevant fixed orientation of the DNA (Fig. 3A) show that the binding of *IκB* not only changes the DNA binding free energy to *NFκB* but also alters the detailed mechanism by which DNA associates with and dissociates from *NFκB*. *IκB* lowers the barrier for DNA escape by at least 2 – 3 *kcal* (black arrows on Fig. 3A). This change implies that a considerable acceleration of the rate of dissociation from the DNA can be induced by binding *IκB*. Looking at the profile from the DNA association viewpoint one sees that there is a significant kinetic barrier for DNA to associate with a preexisting *NFκB* − *IκB* complex, while DNA associates with free *NFκB* in a diffusion limited manner as shown by the purely downhill profile for binary association in the absence of *IκB*.

**Figure 3:**
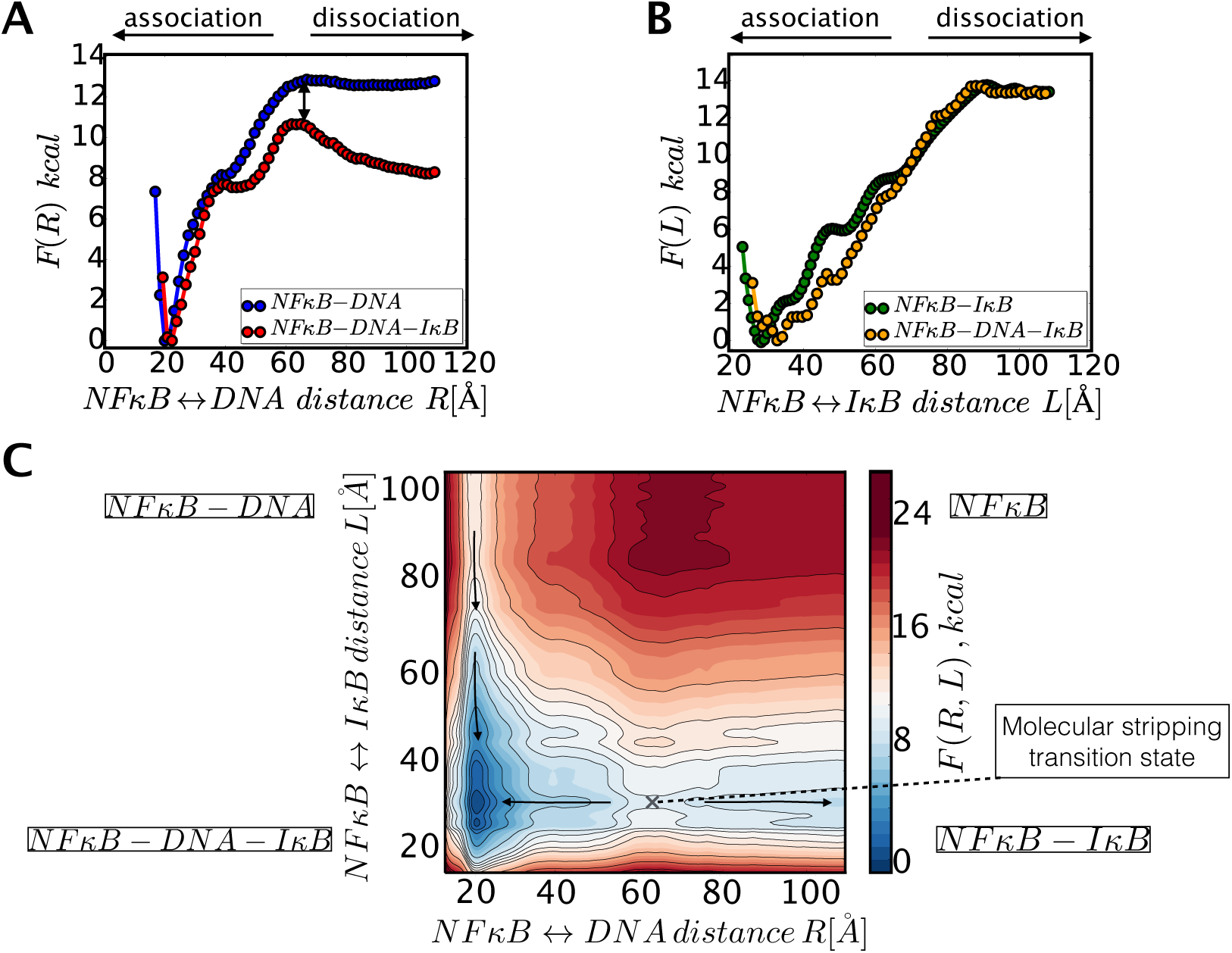
(A) The 1D Free energy profile for *IκB* dissociation from the ternary complex *IκB* − *NFκB* − *DNA* (B) The 1D Free energy profile for DNA dissociation from the binary complex of *NFκB* − *DNA* (in blue) and from the ternary complex of *IκB* − *NFκB* − *DNA* (in red) (C) A free energy surface combining the 1D plots in A and B panels. The initial, intermediate and final configurations on a molecular stripping pathway can be seen in Fig. 6.

Examining the free energy profile for *IκB* dissociation from the ternary complex (Fig. 3B), shows that there is a larger barrier for releasing *IκB* from the complex than for releasing *DNA*. This is in harmony with kinetic mixing experiments that do not observe dissociation of the *IκB* on the laboratory time scale (*7, 12*). Combining the two 1D Free energy profiles into a two dimensional free energy surface (Fig. 3C) maps out the interchange of the two binding partners, through the approach of IκB to the binary complex with DNA and the subsequent dissociation of the DNA from the resulting ternary complex. The dynamical and thermodynamic aspects of molecular stripping can be studied at a higher level of structural detail by using additional collective coordinates as discussed below. The lower barrier for DNA dissociation from the ternary complex *NFκ* − *DNA* − *IκB* compared to binary complex *NFκB* − *DNA* and an even higher barrier for *IκB* dissociation (Fig. 3C) shows that molecular stripping is not only thermodynamically but also kinetically favorable.

## Essential modes of motion

Being a multi-domain protein complex, as expected, the *NFκB* hetero-dimer can bend around the connecting hinge while the individual domains themselves remain relatively rigid. To characterize this and other global slow time-scale motions we performed many exhaustive simulations with a cumulative run time equivalent to more than 500 microseconds of real time. The long sampling times are made possible by using our coarse grained model. Using long constant temperature simulations we first carried out principal component analysis in the Cartesian space spanned by the backbone *C_α_* atoms of the free *NFκB* alone to uncover the dominant large amplitude modes of motion. Principal component analysis of well sampled trajectories of proteins in their native states provides a convenient simplifying lens to characterize their functional collective motions (*31, 34*).

The first principal component is the predominant motion which accounts for more than 70% of the amplitude of motion in unbound *NFκB* (Fig. S4), but a second component also plays a role. Projection of the equilibrium ensemble of structures onto these two modes reveals that these large amplitude motions are not quasi-harmonic in the free *NFκB* dimers, leading to multi-peak distributions for these collective collective coordinates in the free form (Fig. 4A), S5). The first principal component is largely a domain twist motion (Fig. S6, Movie S1). The second principal component for the free *NFκB* is an overall domain breathing motion, resembling the opening and closing of the DNA binding cavity (Fig. S6, Movie S2). In contrast to the free *NFκB* case, for both binary complexes the distribution of these coordinates is essentially single-peaked, with peaks in different locations for *NFκB*−IκB and for *NFκB*−*DNA* (Fig. 4A). *IκB* binding and *DNA* binding couple to distinct motions of the free *NFκB*, namely dominantly to the twist when forming *IκB* − *NFκB* but to the breathing mode when forming *NFκB* − *DNA*.

**Figure 4:**
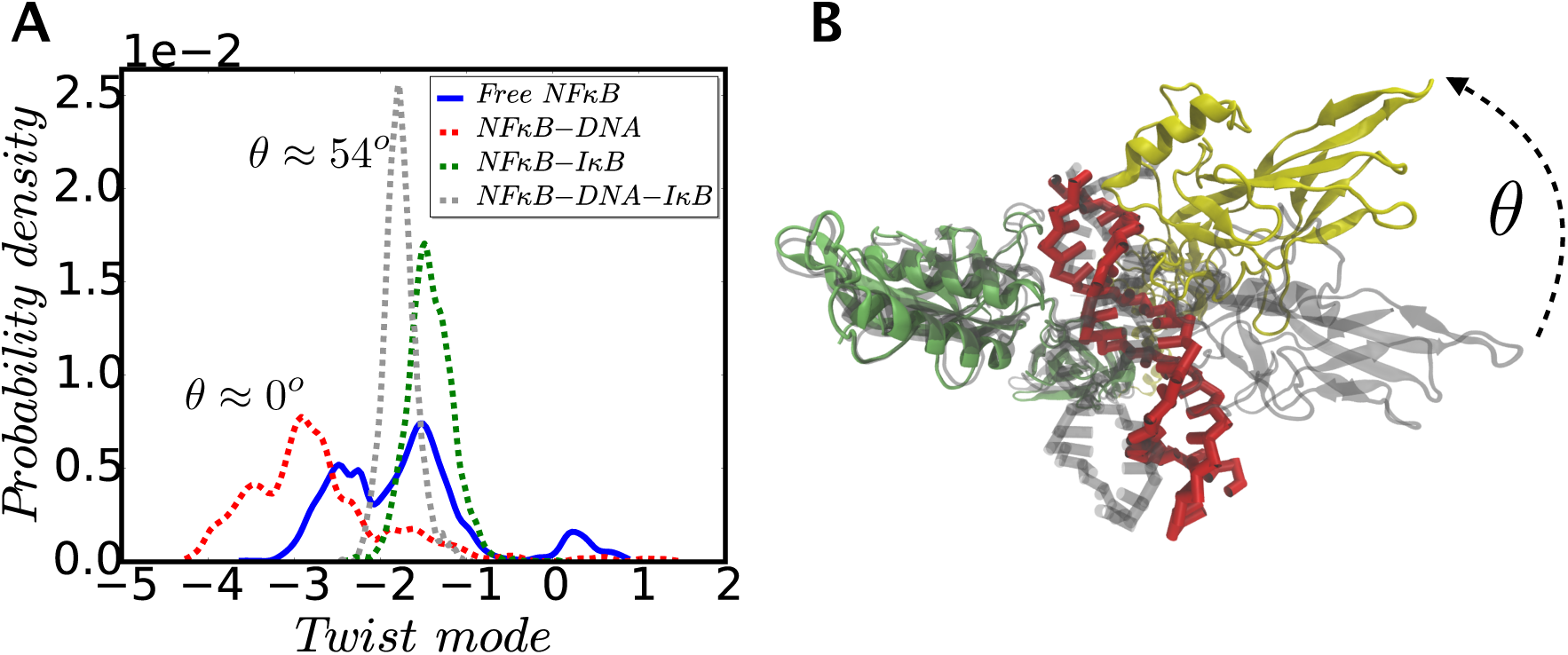
(A) Distribution of conformational ensembles for free and bound forms of *NFκB* along the twist mode coordinate of free *NFκB*. (B) The p50 domain of the binary complex (grey) was aligned with the p50 domain of the ternary *NFκB* − *DNA* − *IκB* complex (p50: green, p65: yellow, DNA: red, *IκB*: not shown). The *IκB* induces a twist in the relative orientation of the p50 and p65 domains (angle *θ*).

Projecting the equilibrium ensemble of the binary complexes on principal components determined now by including both backbone atoms of *NFκB* and the backbone atoms of its binding partners (either DNA or *IκB*), reveals the motions of the partners with respect to each other. The principal motion of the *NFκB* − *DNA* complex corresponds to sliding the *NFκB* along the DNA strands (Fig. S6), supporting the view that *NFκB* can easily perform a 1D diffusional motion along the DNA without being released. The second largest principal component for *NFκB* − *DNA* as a whole complex corresponds to what was the domain twist motion of the free *NFκB*. The domain breathing motion which is displaced upon DNA binding is heavily suppressed in fluctuating amplitude after the DNA binds.

When *IκB* binds to free *NFκB* the induced twist orients the domains so that only one of the *NFκB* monomers can now contact the DNA very closely (Fig. 4B). This observation is one of the key results of this work. We see that *IκB* acts as an allosteric effector precisely by selecting a conformation in the *NFκB* − *DNA* most favorable for DNA dissociation. Once the *IκB* binds the *NFκB*, the *DNA* remains held by only a single N-terminal domain of *NFκB* coming from the p50 subunit (Fig. 4B). In contrast both N-terminal domains contribute to holding strongly to the *DNA* before the twist transition has taken place. The allosteric change through twisting ultimately allows the stripping of *NFκB from its genomic binding* sites on the *DNA*.

## Mechanistic picture of molecular stripping

By mapping the free energy profile simultaneously onto the dissociation distances (Fig. 3) and onto the principal components of free *NFκB* we can uncover the key mechanistic steps by which DNA dissociates from *NFκB* after the complex with *IκB* forms. Following how the domain twist coordinate changes as the partners approach each other we see that the minimal free energy paths for dissociation of DNA have two stages (Fig. 5). In the first stage the domains twist (note the solid arrows on Fig. 5AB), thereby loosening the steric and electrostatic grip of the transcription factor on the *DNA*. This twist is then followed by a subsequent step in which the *DNA* separates from the complex while only one of the two domains holds on strongly to the DNA. The free energy surface for DNA dissociation from the binary complex *NFκB* − *DNA* (Fig. 5 C) shows a bifurcation: the DNA can be induced to dissociate by rotating either one of the two DNA binding domains. The two dissociating paths do not have equal barriers which highlights the symmetry breaking in the hetero-dimer of *NFκB*.

**Figure 5:**
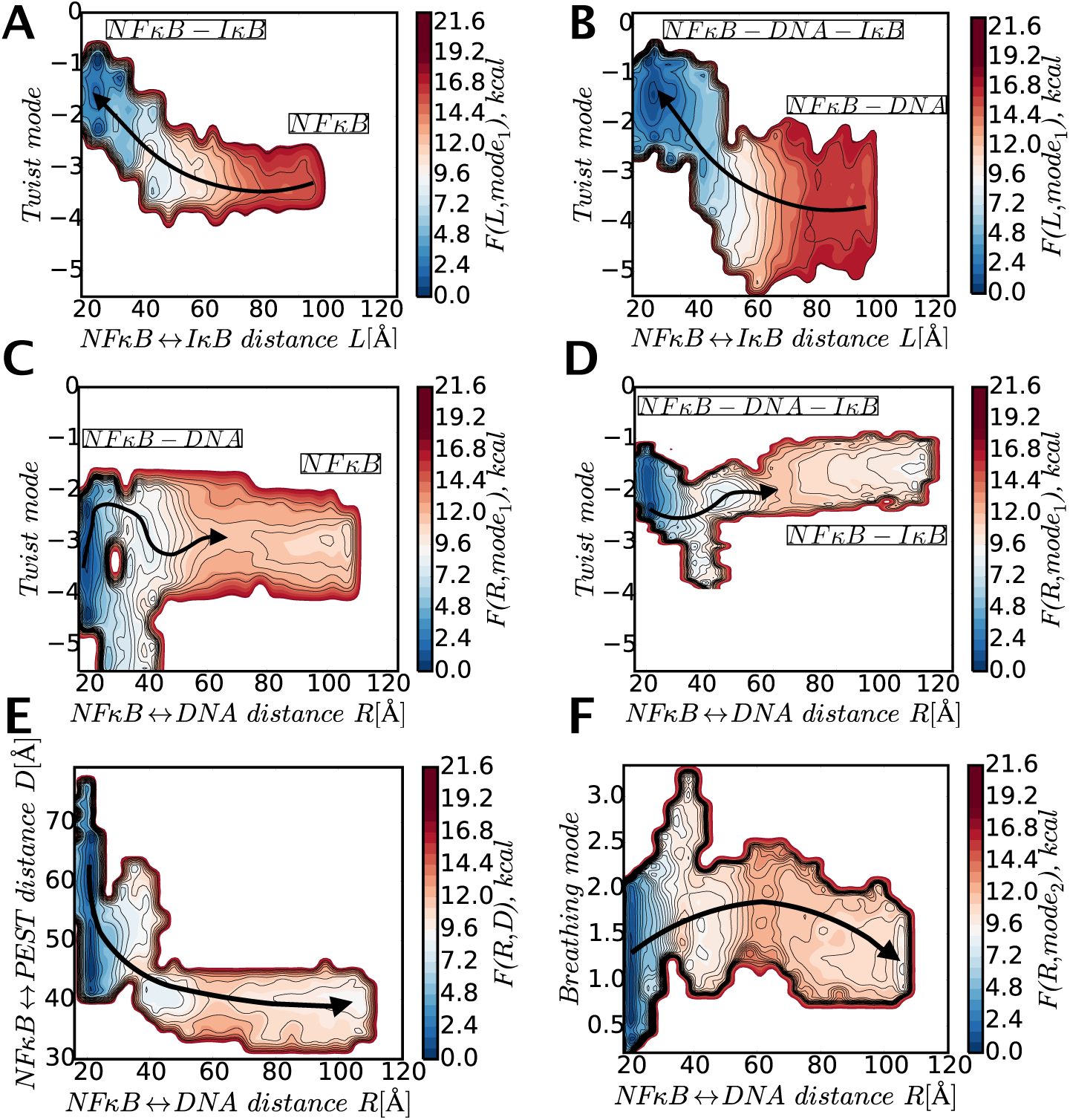
Panels (A-D): Free energy surfaces mapped onto relevant dissociation/association distances and the domain twist mode. Dark arrows indicate the the direction of the most favorable path leading to formation of stable complexes. (A) Free energy surface for *NFκB* binding to *IκB* forming a binary complex *IκB* − *NFκB* (B) Free energy surface for *NFκB − DNA* binding to *IκB* forming a ternary complex *NFκB − DNA − IκB* (C) Free energy surface for DNA dissociation from the binary complex *NFκB − DNA* to form free *NFκB* (D) Free energy surface for DNA dissociation from the ternary complex *NFκB − DNA − IκB* to form the binary complex *NFκB − IκB* (E) Free energy surface mapped onto the DNA dissociation distance and the PEST distance from the binding pocket of the *NFκB* in the ternary complex of the *NFκB − DNA − IκB*. (F) Free energy for dissociation of the DNA − *NFκB* complex mapped onto the DNA dissociation distance and onto the breathing mode that opens and closes the DNA binding cavity.

Dissociation of *DNA* from the ternary complex proceeds by a different route (Fig. 5D and 6) than does *DNA* dissociation from the binary complex because the binding of the *IκB* to the *NFκB* − *DNA* binary complex has already induced the necessary domain twist of the p65 domain via a conformational selection mechanism (Fig. 5B). In forming the ternary complex the *DNA* binding domain has already come to a more favorable orientation for releasing *DNA*, leading to lower kinetic barriers (Fig. 5 D). We see the first step of molecular stripping occurs through an allosteric transition caused by the *IκB* acting on the domain twist mode of the *NFκB* hetero-dimer. This conformational change facilitates the stripping of the *NFκB* from DNA by decreasing the barrier for further domain twist.

**Figure 6:**
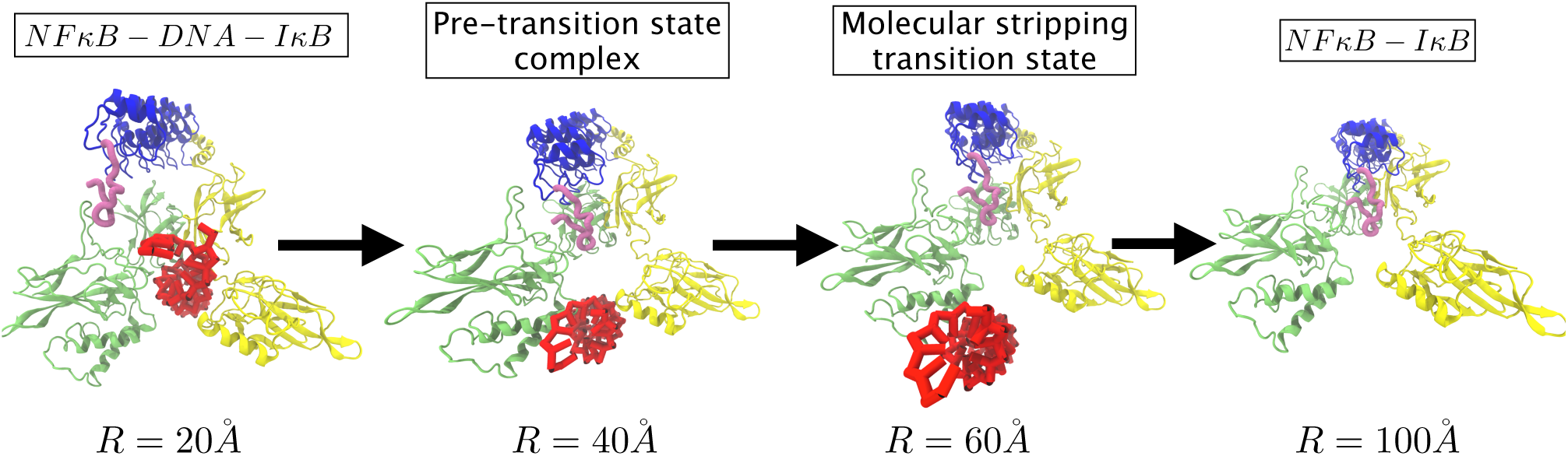
The progression of structural changes on the molecular shipping path starting from the ternary complex *DNA* − *NFκB* − *IκB* and going through the states of the pre-transition state complex, the molecular stripping transition state and ending at the stable binary complex of *IκB*−*NFκB*. The pink tube highlight shows the PEST sequence and its motion with respect to the *NFκB* throughout the DNA dissociation. The partial collapse of the binding cavity once the DNA has left is also evident.

Once DNA leaves the ternary intermediate, the resulting bound complex of *NFκB* − *IκB* is thermodynamically much more stable (Fig. 5A) than the starting complex with *NFκB* bound only to the DNA (Fig. 5B). It is then no surprise that the reverse reaction of *DNA* stripping the *IκB* moiety from its complex with *NFκB* has not been observed in kinetic experiments (*7*). The thermodynamic stability of the *NFκB* − *IκB* complex as well as the origin of barrier for the reverse reaction of DNA binding can be explained using electrostatic arguments as follows. In the second step of stripping, as the DNA begins to leave, the *IκB* reconfigures through the motion of the *PEST* sequence. By being highly negatively charged the PEST sequence begins to compete with the DNA for binding to the positively charged *NFκB* binding pocket (Fig. 5E). This exchange of partners lowers the barrier for DNA dissociation.

As is seen on the free energy surface, once the DNA has completely left the cavity, the *PEST* sequence moves still further to occupy the cavity thereby finally resolving the electrostatically frustrated interactions (Fig. 5E and 6). Looked at hypothetically in the reverse direction of association, the occupation of the DNA binding pocket by the PEST sequence leads to a free energy barrier inhibiting association of the DNA with the *NFκB* − *IκB* complex. This barrier gives another way by which *IκB* controls the transcriptional ability of the *NFκB*. Note that in both the pre-transition state complex and the molecular stripping transition state itself the DNA remains bound transiently to the p50 subunit for which it has the higher affinity (*35*).

One may use other collective modes for investigating different mechanistic aspects of *NFκB* domain motions. For instance using the breathing mode as a collective coordinate (Fig. 5F) reveals a displacement in the breathing mode coordinate when DNA dissociates or binds to the free *NFκB*. The two DNA binding domains of *NFκB* first open and then close, approaching each other to seal the cavity between them thus locking onto the target *DNA*.

## Concluding remarks

Single cell experiments on *NFκB* regulation *in vivo*, mathematical modeling, *in vitro* kinetics and structural studies have provided many important clues to *NFκB* signalling systems biology. The present simulations show that a detailed molecular level understanding of the functional dynamics of the key *NFκB* − *IκB* − *DNA* genetic switch elements can add more to the story. In this work exhaustive long time-scale coarse grained simulations for the free *NFκB* hetero-dimer and its bound complexes with DNA and *IκB* have allowed us to explore structures, thermodynamics and motions that have not been accessible to direct crystallographic study. Our simulations demonstrate that the *NFκB/DNA/IκB* switch functions via an allosteric mechanism which involves conformational selection of a collective mode of domain twist accompanied by opening and closing of the cavity between the two DNA binding domains of the hetero-dimer. The molecular stripping process of *NFκB* from its target genetic sites by *IκB* proves both kinetically and thermodynamically favorable suggesting a novel non-equilibrium mechanism for the efficient control of gene expression in this important master regulator system.

## References and Notes

1. M. Ptashne, Nature 322, 697 (1985).

2. M. Ptashne, A genetic switch: phage lambda revisited, vol. 3 (Cold Spring Harbor Laboratory Press Cold Spring Harbor, NY:, 2004).

3. W. Gilbert, B. Müller-Hill, Proc Natl Acad Sci 56, 1891 (1966).

4. F. Jacob, J. Monod, J Mol Biol 3, 318 (1961).

5. G. K. Ackers, A. D. Johnson, M. A. Shea, Proc Natl Acad Sci 79, 1129 (1982).

6. R. Phillips, J. Kondev, J. Theriot, H. Garcia, Physical biology of the cell (Garland Science, 2012).

7. V. Alverdi, B. Hetrick, S. Joseph, E. A. Komives, Proc Natl Acad Sci 111, 225 (2014).

8. A. Oeckinghaus, S. Ghosh, Cold Spring Harb Perspect Biol 1, a000034 (2009).

9. D. Krappmann, et al., Mol Cell Biol 24, 6488 (2004).

10. A. Hoffmann, A. Levchenko, M. L. Scott, D. Baltimore, Science 298, 1241 (2002).

11. D. Nelson, et al., Science 306, 704 (2004).

12. S. Bergqvist, et al., J Mol Biol 360, 421 (2006).

13. P. A. Baeuerle, D. Baltimore, Science 242, 540 (1988).

14. M. D. Jacobs, S. C. Harrison, Cell 95, 749 (1998).

15. R. A. Williams, J. Timmis, E. E. Qwarnstrom, Computation 2, 131 (2014).

16. R. Cheong, A. Hoffmann, A. Levchenko, Mol Sys Biol 4, 192 (2008).

17. S. Bergqvist, et al., Proc Natl Acad Sci 106, 19328 (2009).

18. R. Martone, et al., Proc Natl Acad Sci 100, 12247 (2003).

19. B. Berkowitz, D.-B. Huang, F. E. Chen-Park, P. B. Sigler, G. Ghosh, J Biol Chem 277, 24694 (2002).

20. T. Huxford, D.-b. Huang, S. Malek, G. Ghosh, Cell 95, 759 (1998).

21. A. Davtyan, et al., J Phys Chem B 116, 8494 (2012).

22. D. M. Hinckley, G. S. Freeman, J. K. Whitmer, J. J. de Pablo, J Chem Phys 139, 144903 (2013).

23. G. S. Freeman, D. M. Hinckley, J. P. Lequieu, J. K. Whitmer, J. J. de Pablo, J Chem Phys 141 (2014).

24. J. N. Onuchic, Z. Luthey-Schulten, P. G. Wolynes, Ann Rev Phys Chem 48, 545 (1997).

25. W. Zheng, N. P. Schafer, A. Davtyan, G. A. Papoian, P. G. Wolynes, Proc Natl Acad Sci 109, 19244 (2012).

26. D. A. Potoyan, A. Savelyev, G. A. Papoian, WIRES Comp Mol Sci 3, 69 (2013).

27. J. J. de Pablo, Annu Rev Phys Chem 62, 555 (2011).

28. J. D. Bryngelson, P. G. Wolynes, Proc Natl Acad Sci 84, 7524 (1987).

29. J. D. Bryngelson, J. N. Onuchic, N. D. Socci, P. G. Wolynes, Proteins 21, 167 (1995).

30. D. U. Ferreiro, E. A. Komives, P. G. Wolynes, Quart Rev Biophys 47, 285 (2014).

31. P. I. Zhuravlev, G. A. Papoian, Quart Rev Biophys 43, 295 (2010).

32. M. Jenik, et al., Nucl Acid Res p. 447 (2012).

33. M.-Y. Tsai, et al., Protein Science (2015).

34. D. A. Potoyan, G. A. Papoian, J Am Chem Soc 133, 7405 (2011).

35. C. B. Phelps, L. L. Sengchanthalangsy, S. Malek, G. Ghosh, J Biol Chem 275, 24392 (2000).

36. We thank Holly Dembinsky, Kristen Ramsey, Bin Zhang and Patricio Craig for numerous stimulating discussions. This work is supported by PPG Grant P01 GM071862 from the National Institute of General Medical Sciences. P.G.W. also gratefully acknowledges financial support from the D.R. Bullard-Welch Chair at Rice University, Grant C-0016.

